# Monosomies, trisomies and segmental aneuploidies differentially affect chromosomal stability

**DOI:** 10.1101/2021.08.31.458318

**Authors:** Dorine C. Hintzen, Mar Soto, Michael Schubert, Bjorn Bakker, Diana C.J. Spierings, Karoly Szuhai, Peter M. Lansdorp, Floris Foijer, René H. Medema, Jonne A. Raaijmakers

## Abstract

Aneuploidy and chromosomal instability are both commonly found in cancer. Chromosomal instability leads to karyotype heterogeneity in tumors and is associated with therapy resistance, metastasis and poor prognosis. It has been hypothesized that aneuploidy *per se* is sufficient to drive CIN, however due to limited models and heterogenous results, it has remained controversial which aspects of aneuploidy can drive CIN. In this study we systematically tested the impact of different types of aneuploidies on the induction of CIN. We generated a plethora of isogenic aneuploid clones harboring whole chromosome or segmental aneuploidies in human p53-deficient RPE-1 cells. We observed increased segregation errors in cells harboring trisomies that strongly correlated to the number of gained genes. Strikingly, we found that clones harboring only monosomies do not induce a CIN phenotype. Finally, we found that an initial chromosome breakage event and subsequent fusion can instigate breakage-fusion-bridge cycles in segmental aneuploidies. This suggests that monosomies, trisomies and segmental aneuploidies have fundamentally different effects on chromosomal instability and these results help us to decipher the complex relationship between aneuploidy and CIN.

## Introduction

Chromosomal instability (CIN) and aneuploidy are both acknowledged hallmarks of cancer. Although these terms are often used interchangeably, they have different implications and it is therefore crucial to stress the difference. While aneuploidy refers to the genomic status of a cell with an abnormal chromosome content at a given time, CIN refers to the behavior of cells that display dynamic chromosomal content due to persistent segregation errors (Lengauer et al., 1998). There are two main types of aneuploidy; numerical aneuploidy, referring to the gain or loss of whole chromosomes, and segmental aneuploidy, referring to the gain or loss of parts of chromosomes. Besides gaining or losing genetic information, segmental aneuploidies also often result in structural chromosomal aberrations such as translocations and dicentric chromosome formation (Soto et al., 2019).

Aneuploidy has been shown to be detrimental in all organisms, leading to growth and developmental impairments (Santaguida and Amon, 2015; Torres et al., 2008). However, aneuploidy is very common in cancer with approximately 90% of all solid tumors harboring abnormal karyotypes (Taylor et al., 2018). Single cell techniques have shown that besides stable aneuploidies, cells often display karyotype variation within a single tumor, due to ongoing CIN. Ongoing CIN allows cells to widely sample a range of different genotypes that can eventually be selected to match the requirements faced under stress (Chen et al., 2012). Indeed, continuous karyotype deviations and subsequent karyotype selection have been shown to confer adaptation to challenging environments in single-cell organisms (Chang et al., 2013; Pavelka et al., 2010; Selmecki et al., 2006; Yona et al., 2012) as well as in mammalian cell culture (Ippolito et al., 2021; Lukow et al., 2021; Rutledge et al., 2016). Moreover, several studies have demonstrated that ongoing segregation errors in tumors are associated with enhanced therapy resistance, poor prognosis, and metastasis, thus reflecting adaptation conferred by CIN (Bakhoum et al., 2018; Swanton et al., 2009). However, the oncogenic potential of CIN can differ per tissue and high levels can in some cases also decrease tumorigenic potential (Birkbak et al., 2011; Janssen and Medema, 2012; Weaver et al., 2007; Hoevenaars et al. 2020).

Due to the high prevalence of aneuploidy and CIN in cancer, much attention has been put over the last years to reveal the underlying causes. Some causes have been identified such as a weakened spindle checkpoint, defects in chromosome cohesion, abnormal kinetochore-microtubule attachments and supernumerary centrosomes (Sansregret and Swanton, 2017; Bastians, 2015; Thompson et al., 2010). Nevertheless, mutations in key regulators of these processes are rarely found in tumors (Sansregret and Swanton, 2017), and thus the exact underlying cause for CIN remains unclear in the majority of cases. While aneuploidy is a consequence of CIN, it remains controversial if aneuploidy *per se* can be an underlying cause of CIN. Some studies have indeed suggested that deviating karyotypes can instigate instability (Sheltzer et al., 2011; Zhu et al., 2012; Duesberg et al., 1998; Nicholson et al., 2015; Passerini et al., 2016; Thompson and Compton, 2010). For instance, CIN has been suggested to be the result of specific gains or losses of chromosomes that contain key regulators of proper segregation in mitosis (Nicholson et al., 2015; Zhu et al., 2012). Furthermore, it has been shown that the presence of extra chromosomes disrupts proteostasis and saturation of the protein folding machinery can result in misfolded proteins, aggregate formation and activation of protein degradation pathways (Dürrbaum et al., 2014; Stingele et al., 2012, 2013; Tang et al., 2011). Subsequently, this can result in the deregulation of specific proteins that highly depend on the chaperone machinery. These proteins involve, among others, key players of replication, therefore expression changes possibly increase replication stress and lead to CIN (Passerini et al., 2016). Importantly, it has also been shown that not all whole chromosomes imbalances induce CIN (Lengauer et al., 1997; Sheltzer et al., 2011; Zhu et al., 2012), indicating that the mere presence of additional chromosomes may not always be sufficient to induce chromosomal instability.

Here, we set out to investigate the impact of a large panel of *de novo* induced stable aneuploidies on chromosome stability in a near-diploid human cell line. Specifically, we aim to understand whether different types of aneuploidies have a different impact on CIN, as well as understand what features of aneuploidy drive instability. Importantly, while most studies have focused mostly on trisomies, in this study we also include aneuploid clones with only monosomies, segmental aneuploidies as well as more complex karyotypes.

## Results

### Generation of 24 isogenic aneuploid clones

To study the effects of aneuploidy on chromosome instability, we generated a large panel of aneuploid clones derived from an hTERT-immortalized retinal pigment epithelial cell line (RPE-1). As p53 has an important role in aneuploidy tolerance (Santaguida et al., 2017; Soto et al., 2017; Thompson and Compton, 2010) and is mutated in the majority of cancers, aneuploid clones were generated under a stable shRNA knockdown of p53 (Soto et al., 2017). For this, a total of ten 384-well plates (3840 wells) in 5 independent experiments were seeded with single cells that were pretreated overnight with a low dose of MPS1 and CENP-E inhibitors to induce chromosome missegregations. Each well was subsequently analyzed and approximately 650 wells contained a single cell by visual inspection. About ∼6% of these cells made it to a full clone and only a subset of those had a confirmed aneuploid karyotype. With this approach, we obtained a total of 24 clones containing one or more *de novo* aneuploidies (since the gain of chromosome 10q and the gain of chromosome 12 are common events in parental cells, these were excluded as *de novo* events; for more details see Materials and Methods). A subset of clones contained solely whole chromosome imbalances (Fig. S1), whereas others contained segmental abnormalities, sometimes in combination with whole chromosome aneuploidies (Fig. S2). Segmental aneuploidies have undergone DNA breaks and potentially aberrant DNA repair resulting in abnormal chromosomes. As this could complicate the analysis of the sole effect of aneuploidy on CIN, we first focused on clones exclusively harboring whole chromosome aneuploidies with clean CNV profiles (15 clones, Fig. S1), as these clones are less likely to have experienced chromosome damage during their generation. As a control, we selected two single cell derived clones (C1 and C2) that displayed a karyotype comparable to the parental cell line but underwent the same procedure as the other established aneuploid clones.

### Aneuploid clones show a spectrum of different doubling times and missegregation rates

We set out to characterize our various aneuploid clones. One of the known consequences of aneuploidy is reduced cellular fitness and proliferation (Sheltzer et al., 2012; Torres et al., 2007; Williams et al., 2008). To assess the proliferation rates of our clones, we determined the doubling times with live-cell imaging. As expected, the majority of our aneuploid clones displayed impaired growth compared to the euploid controls (Fig. 1A). The proliferation impairment was heterogeneous between the different clones with doubling times ranging from 16h to 72h. As we aim to shed light on the relationship between whole chromosome aneuploidy and CIN, we evaluated the levels of CIN in our clones by live cell imaging (Fig. 1B). The parental cell line and the two control clones displayed a basal level of segregation errors of ∼7-10%. When assessing the levels of CIN in our clones, we found that aneuploid clones showed a spectrum of missegregation rates, ranging from ∼5-70% (Fig. 1A). Strikingly, not all clones showed an increase in segregation errors when compared to the controls. This observation demonstrates that imbalances of whole chromosomes can trigger a spectrum of CIN levels, however, not all aneuploidies instigate a CIN phenotype.

**Figure 1.**
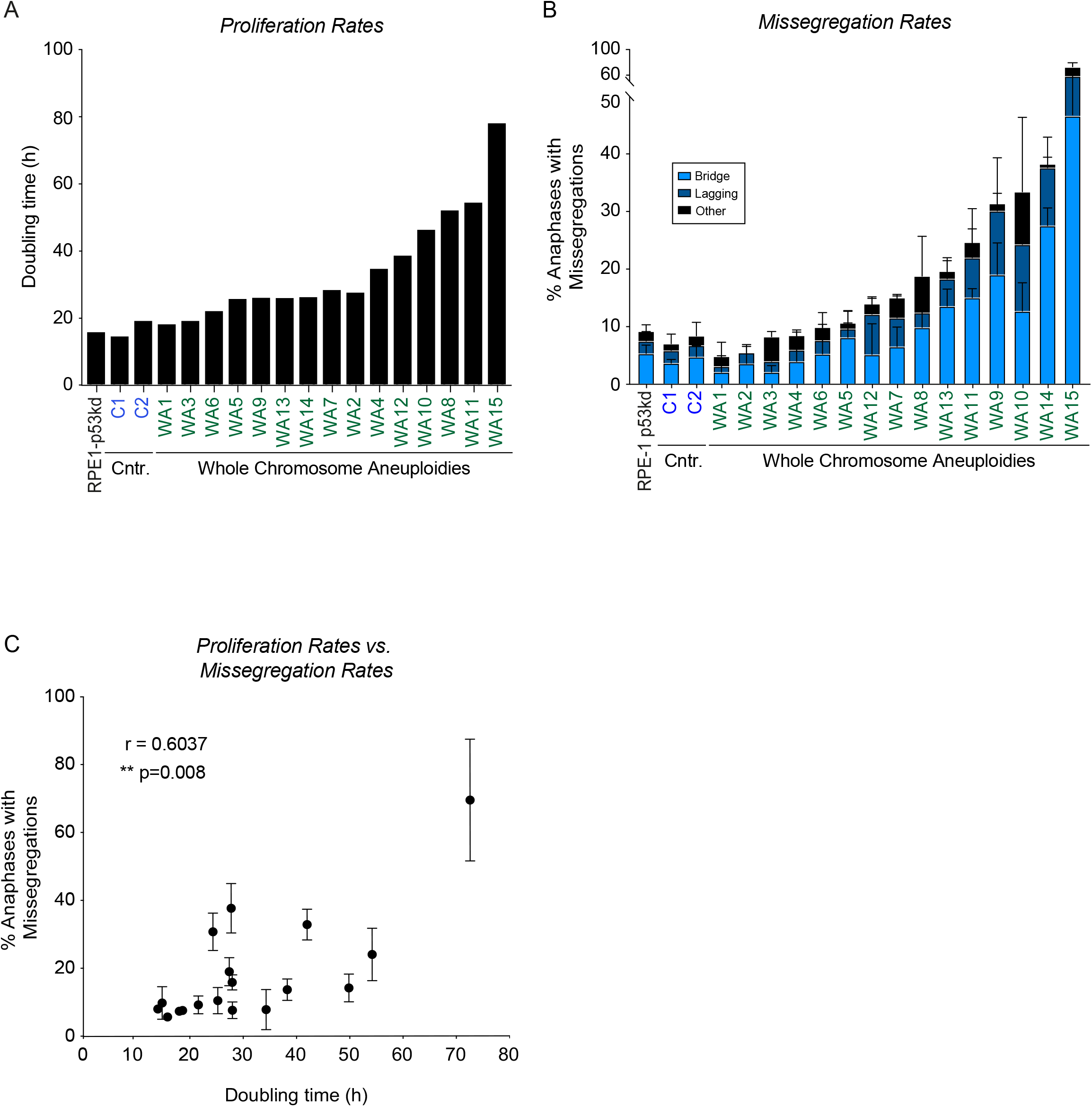
Aneuploid clones show a spectrum of different doubling times and mis-segregation rates. A. Doubling time of the parental cell line (labelled in black), two euploid clones (blue) and aneuploid clones (green). B. Chromosome missegregation rates determined by live cell imaging of parental RPE-1 p53KD cells (labelled in black), euploid (blue) and aneuploid (green) clones from Fig. S1, divided into three subcategories: lagging chromosomes, anaphase bridges and others. Bars are averages of at least 2 experiments and a minimum of 50 cells were filmed per clone. Error bars indicate standard deviation. C. Spearman correlation between the proliferation rate as measured in A and the missegregation rates as a percentage of anaphases as measured in B.

To understand if CIN could be an underlying cause for decreased cellular fitness we evaluated the correlation between doubling times and missegregation rates. Interestingly, we found a significant positive correlation (Fig. 1C, p = 0.008). However, important to note is that some of our chromosomally stable clones display a clear growth impairment (e.g. clone WA2 and WA4, compare Fig. 1A to Fig. 1B), while other highly unstable clones seem to have a milder proliferation defect (e.g. clone WA9 and clone WA14). This suggests that although CIN and proliferation impairment are both consequences of aneuploidy that could be linked to each other, they unlikely have a direct causative relationship.

### Proliferation defects are linked to gene imbalances

Previously, it has been suggested that the impaired proliferation rates in aneuploid yeast cells are largely determined by the increased dosage of coding genes, as additional non-coding DNA did not cause a growth defect (Torres et al., 2007). Importantly, a more severe growth impairment could be observed when larger or more chromosomes were gained (Torres et al., 2007). The effect of chromosome losses has not been addressed as the studied yeast model contains a haploid genome to start with. We set out to determine if the degree of aneuploidy could also explain the variation in proliferation rates observed in our mammalian cell system. We calculated the degree of aneuploidy by determining the total amount of coding genes that are imbalanced per clone. Indeed, we found a positive correlation between doubling times and number of imbalanced genes (Fig. 2A, left panel, p = 0.0007). When separating the imbalanced genes in lost and gained genes we found that proliferation rates were affected by both the gain and the loss of genetic material, as both independently showed a significant correlation to proliferation rates (Fig. 2A, middle and right panel). However, neither the gained genes nor lost genes provided additional information about the growth rate over the number of imbalanced genes, as the number of imbalanced genes could fully explain the two other correlations (conditional Spearman tests p=0.68 and p=0.60, respectively). Therefore, our data suggests that the reduced proliferation rates are most likely a consequence of gene imbalances.

**Figure 2.**
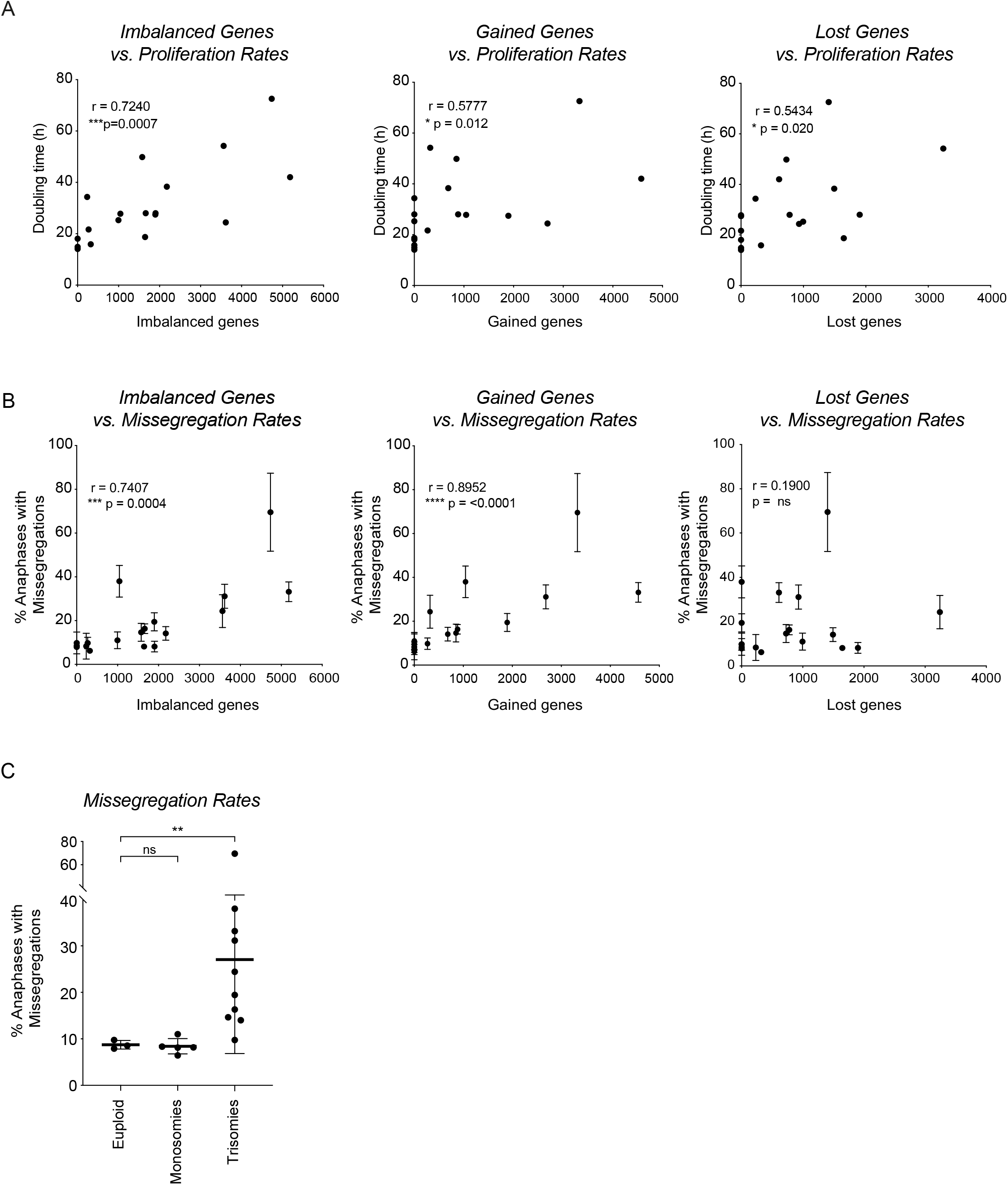
Both monosomies and trisomies decrease cellular fitness. A. Spearman correlation between the total amount of imbalanced, gained and lost coding genes per clone and proliferation rates as measured in Fig. 1A. B. Spearman correlation between the number of imbalanced, gained and lost coding genes and the level of CIN as percentage of total amount of anaphases as determined in Fig. 1B. Error bars indicate standard deviation. C. Missegregation rates described in A, classified in three different categories: euploid clones, clones harboring only monosomies and clones harboring trisomies (sometimes with monosomies in the background). Lines show the mean; error bars indicate standard deviation. The Mann-Whitney U statistical test was used to compare differences in CIN between groups.

### CIN rates are explained by gained genes rather than by imbalances

Next, we investigated the relationship between the degree of aneuploidy and the levels of CIN. Studies in yeast suggested that instability is also a consequence of dosage changes in coding genes as the addition of non-coding DNA does not instigate a CIN phenotype while extra coding DNA does (Sheltzer et al., 2011). It was previously shown that the more cells deviate from their true euploid state, the more unstable they become (Duesberg et al., 1998; Zhu et al., 2012). However, a clear correlation between the amount of gained material and CIN has not been found (Sheltzer et al., 2011). Here, we found a significant correlation between the amount of imbalanced coding genes and missegregations rates in our mammalian cell system (Fig. 2B, left panel, p = 0.0004). Interestingly, and unlike the correlation between doubling times and degree of aneuploidy, this correlation further improved when only tested against the amount of gained coding genes (Fig. 2B, middle panel, p<0.0001). There was no significant correlation between the amount of lost coding genes and CIN levels (Fig. 2B, right panel). Importantly, the correlation between imbalanced genes and CIN rates could be fully explained by the amount of gained genes (conditional Spearman test for imbalanced genes after correction for gained genes is p = 0.50), but the correlation of gained genes with CIN rates remained significant after correcting for the total number of imbalanced genes (conditional Spearman test p=0.001). Together, these data suggest that gaining extra coding genetic material can lead to CIN while losing coding genetic material does not significantly contribute to this phenotype. Moreover, it shows that proliferation rates and CIN rates are two independent features of aneuploidy that have different underlying causes. Indeed, and in line with our previous hypothesis, we could confirm that reduced proliferation rates can be explained by gene imbalances and not by the CIN phenotype (conditional Spearman test p = 0.33).

In line with the absence of a correlation between the loss of genetic material and CIN rates, we found that clones that did not induce instability mostly involved clones with only monosomies, whereas clones with increased instability exclusively involved clones harboring trisomies. Indeed, when classifying clones by their type of karyotype aberration, the monosomic clones (referred to as monosomies) did not display increased chromosomal instability compared to the control cell lines, while clones that harbor at least one trisomy (referred to as trisomies) in most cases induced CIN to different extents (Fig. 2C). This further underlines our hypothesis that extra genetic material can drive CIN while we did not find evidence that loss of genetic material does so.

### Interfering with proteostasis induces CIN

To further investigate the relation between CIN and gained coding genes, we aimed to elucidate a potential causal relationship. It has been extensively shown that gaining extra coding DNA leads to an excess of proteins being expressed from the involved chromosome (Dephoure et al., 2014; Liu et al., 2017; Pavelka et al., 2010; Stingele et al., 2012; Torres et al., 2010). Both chromosome gains and losses lead to specific protein imbalances. However, chromosome gains result in the enhanced expression of many genes simultaneously and can cause a general overburden of the protein folding and degradation pathways in an attempt to maintain proteostasis. We therefore hypothesized that a general stress response associated to protein overexpression could be a direct or indirect underlying cause for the CIN phenotype in trisomic clones and would explain the absence of CIN in monosomies. First, we aimed to investigate if inducing proteotoxic stress would be sufficient to drive CIN. To achieve this, we interfered with proteostasis in parental cells using the chaperone protein Hsp90 inhibitor 17-AAG or the proteasome inhibitor MG132, using low doses that did not prevent cells from entering mitosis (Fig. S3A). We found that cells treated with either one of the inhibitors showed missegregation rates comparable to the ranges we observed in our clones (Fig. 3A). These data suggest that interfering with protein folding or protein degradation in parental cells is sufficient to trigger a CIN phenotype.

**Figure 3.**
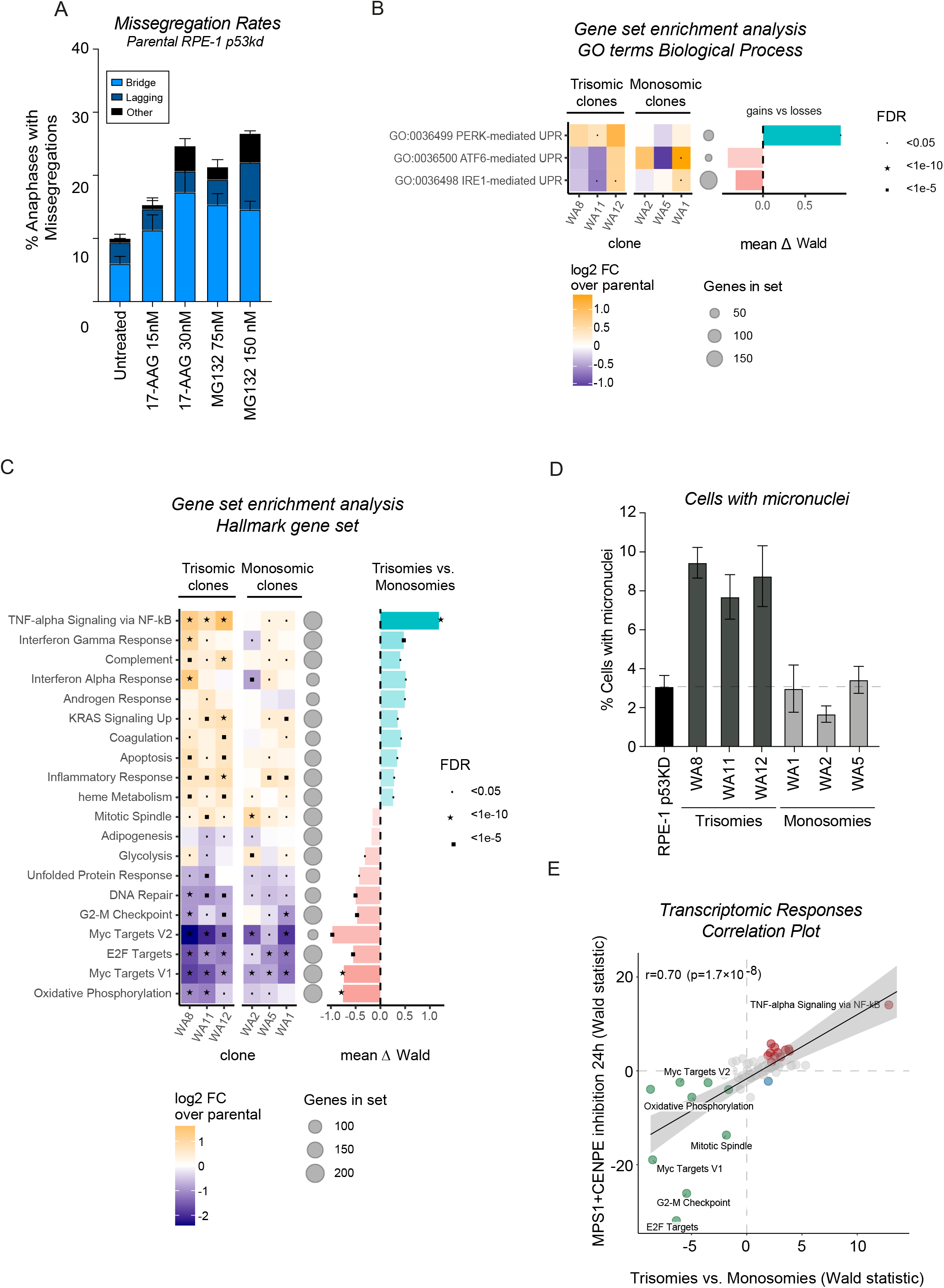
Different transcriptomic responses to trisomies and monosomies. A. Chromosome missegregation rates of RPE-1 parental p53KD cells untreated or treated with low doses of the indicated inhibitors for 24 hours determined as in Fig. 1B. The experiment was performed in triplo and 50 cells were analyzed per condition per experiment. Error bars indicate standard deviation. B. Gene set enrichment analysis (GSEA) specifically for three different UPR branches, evaluating their up and downregulation in trisomic and monosomic clones compared to parental cells (left graph). Two replicates of every clone were sequenced and the Log2 fold change (FC) was determined compared to parental cells. The false discovery rates (FDR) are indicate with symbols. Difference between trisomies and monosomies were determined by Wald statistical testing. C. Gene set enrichment analysis (GSEA) of RNA sequencing data, evaluating up and downregulated Hallmarks in trisomy clones and monosomy clones compared to parental (left graph). Two replicates of every clone were sequenced and the Log2 fold change (FC) was determined compared to parental cells. The false discovery rates (FDR) are indicate with symbols. Largest differences between trisomies and monosomies are shown on the right. Differences in Hallmarks between trisomies and monosomies were determined by Wald statistical testing. D. Percentage of cells harboring micronuclei, as determined via snapshots from live-cell imaging data. 2 experiments were analyzed per clone, a minimum of 150 cells was analyzed per clone per experiment. Bars show the average of 2 experiments; error bars indicate standard deviation. E. Correlation between upregulated and downregulated Hallmarks in acute aneuploidy and the upregulated and downregulated Hallmarks showing the largest difference between trisomies and monosomies determined by Wald statistical testing.

As chemically interfering with proteostasis is sufficient to drive CIN, we were wondering if our trisomic clones indeed experience proteotoxic stress that could explain their CIN. To investigate this, we first evaluated autophagic flux, by examining the conversion between LC3B-I to LC3B-II on autophagosomes (Kabeya et al., 2000; Mizushima et al., 2004). This conversion indicates an activated autophagy pathway which aids in the degradation of misfolded/unfolded proteins upon proteotoxic stress and has previously been shown to be upregulated in trisomic cell lines and in cells where aneuploidy was induced with inhibitors (Santaguida et al., 2015; Stingele et al., 2012). We tested 3 of our trisomic clones that displayed a variety of CIN levels but we could not detect a significant increase in LC3B-II in these clones (Fig. S3C). It needs to be noted that we could also not observe increased LC3B-II levels in parental RPE-1 cells treated with low doses of proteostasis inhibitors that were sufficient to induce CIN (Fig. S3B), suggesting that this assay might not be sensitive enough to detect low levels of proteotoxic stress or that our clones in fact do not increase autophagic flux efficiently. As a second effort, we set out to test our cells for sensitivity to the Hsp90 inhibitor 17-AAG. A previous study has shown that trisomic MEFs are more sensitive to this compound (Tang et al., 2011). Interestingly, the clone with the most trisomies (clone WA10, trisomic for 5 chromosomes) displayed the highest sensitivity to this compound (Fig. S3D). However, although the majority of tested trisomic clones showed slightly higher sensitivity, some clones showed a similar sensitivity or were even more resistant compared to parental cells and two of the monosomic clones displayed enhanced sensitivity compared to parental cells (Fig. S3D). Importantly, the amount of gained genes did not correlate with the sensitivity to the drug (R=-0.3, p=n.s.) and neither did the number of imbalanced genes (R=0.06, p=n.s.). Taken together, interfering with proteostasis in parental cells induces similar CIN levels as observed in our trisomic clones. However, two directed experiments could not provide strong evidence that proteotoxic stress (at least activation of the autophagy pathway or sensitivity to protein folding inhibitors) is consistently present in these clones. It is possible that the levels of proteotoxic stress are below detection limit, or that we assessed aspects of proteostasis that are not relevant to the CIN phenotype.

### Trisomic clones show preferential activation of PERK-mediated Unfolded Protein Response

As we could not directly show the presence of proteotoxic stress in our trisomic clones by two directed assays, we decided to perform RNA sequencing as an unbiased approach to find evidence for proteotoxic stress in trisomic cells. Moreover, with this approach we can identify the most prominent differences between monosomic and trisomic clones that could attribute to the induced CIN in trisomic clones. For this we selected 3 monosomic clones (WA1, WA2 and WA5) and 3 trisomic clones (WA8, WA11 and WA12). We analyzed differential expression compared to parental cells for each clone and determined the most significant genes and gene sets between the two subcategories (trisomies versus monosomies). The most relevant Hallmark for proteotoxic stress is the unfolded protein response (UPR), which encompasses a transcriptional and translational response to endoplasmic reticulum (ER) stress resulting in repressed translation and apoptosis depending on the extent and duration of the response. Unexpectedly, we found that the majority of aneuploid clones in fact displayed a downregulated UPR Hallmark compared to the parental cells (Fig. 3B). As the UPR consists of three main branches; PERK, ATF6 and IRE1 pathways, we decided to evaluate each pathway separately. This analysis showed that the PERK branch of the UPR was significantly upregulated specifically in our trisomic clones, whilst the other branches showed inconsistent results between trisomies and monosomies (Fig. S4A). This indicates that ER-stress might indeed be activated in trisomic clones and not in monosomic clones and this specifically involves the PERK/ATF4-branch. Taken together, we find 1) a strong correlation between gained genes and CIN, 2) that interfering with proteostasis is sufficient to induce CIN and 3) that the PERK pathway is specifically upregulated in trisomic clones. Potentially, this indicates that ER-stress can be linked to CIN.

### Trisomic clones show CIN-mediated differential expression

Besides focusing on the differential regulation of ER stress, we decided to analyze our data to find the pathways that were affected most prominently in trisomic clones as compared to monosomic clones to potentially reveal additional links between chromosome gains and CIN. We found that several cell cycle-related Hallmarks (E2F targets, Myc targets, G2-M checkpoint) were downregulated in all our aneuploid clones but more prominently in the trisomic clones (Fig. 3C), which agrees with the decreased proliferation rates found in these clones. Interestingly, we found that the trisomic clones showed upregulation of several Hallmarks associated with an inflammatory response, such as TNF-alpha signaling via NF-kB, interferon responses, complement and inflammatory response. These responses were either absent, downregulated or only mildly activated in the monosomic clones (Fig. 3B). Rather than a cause for CIN, the pronounced upregulation of inflammatory responses could be a consequence of the elevated levels of CIN for instance via micronuclei formation and activation of the cGAS-STING pathway (Dou et al., 2017; MacKenzie et al., 2017). Indeed, and in line with our missegregation data, we observed an increase in micronuclei in clones harboring trisomies but not in our monosomic clones (Fig. 3C). These data suggest that the elevated CIN could be responsible for driving parts of the transcriptional responses in trisomic clones. To understand which transcriptomic responses in trisomies are explained by the enhanced CIN levels, we performed RNA sequencing of parental cells treated with a low dose of Mps1 and CENP-E inhibitors for 24 hours to induce acute chromosomal instability. We next investigated which Hallmarks that were preferentially affected in trisomic clones correlated to the Hallmarks affected in parental cells with acute CIN, to determine the responses that are likely driven by CIN. Indeed, we found an overall significant correlation between the Hallmarks that were preferentially affected in trisomic clones compared to parental cells with acute CIN (Fig. 3D). Most prominently, we found a very strong upregulation for TNF-alpha signaling via NF-kB. Thus, the inflammatory response that is unique to trisomies is indeed likely explained by the fact that these clones are more CIN.

In order to get a more fine-grained understanding of the transcriptional changes between trisomies and monosomies, we also quantified which Gene Ontology (GO) Biological Process categories were expressed at different levels. We found that pathways associated with translation and ribosome biogenesis were downregulated in all clones (rRNA processing, translation initiation) (Fig. S4B). Although in monosomies this downregulation was previously attributed to haploinsufficiency of ribosomal genes (Chunduri et al., 2021), our data suggest that chromosome gains can possibly also lead to deregulation of these pathways, possibly due to a general aneuploidy-induced stress response (Terhorst et al., 2020). Taken together, although trisomic cells and monosomic cells display different transcriptional responses, the majority of these differences can be explained by their CIN phenotype.

### Segmental chromosome aneuploidies can induce CIN via BFB-cycles

Besides clones with whole chromosome aneuploidies, we also obtained clones carrying segmental aneuploidies with or without additional whole chromosome imbalances (12 clones, Fig. S2). When analyzing CIN levels in these clones, we again observed that not all segmental aneuploidies instigate a CIN phenotype (e.g. clones SA5 and SA7) (Fig. 4A). Consistently with our clones harboring whole chromosome monosomies, clone SA5 and SA7 harbor solely segmental losses (Fig. S2). Moreover, we could observe that a spectrum of instabilities was induced by the clones harboring segmental aneuploidies (Fig. 4A, B). Consistently, we observed a strong correlation between the number of gained genes and CIN rates (Fig. 4C, middle panel, p= 0.0015). Again, there was no significant correlation between lost genes and CIN levels, as observed in clones harboring whole chromosome aneuploidies (Fig. 4C, right panel)

**Figure 4.**
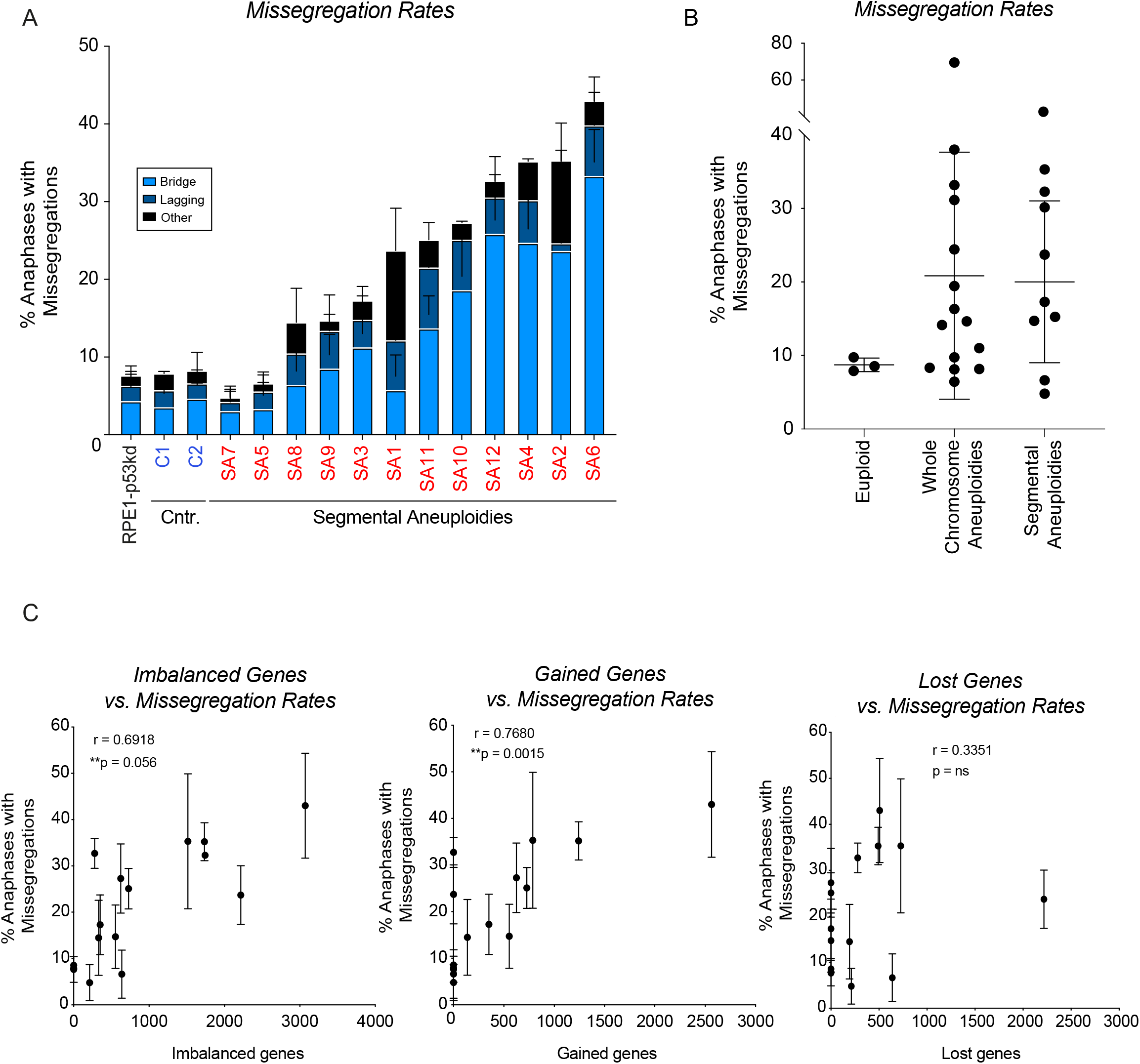
Segmental chromosome aneuploidies can induce CIN. A. Chromosome missegregation rates determined by live cell imaging of parental RPE-1 p53KD cells (labeled in black), euploid clones (blue) from Fig S1 and the segmental aneuploid clones (red) from Fig. S6, categorized in three different categories: lagging chromosomes, anaphase bridges and others. Bars are averages of at least 2 experiments and a minimum of 50 cells were filmed per clones. Error bars indicate standard deviation. B. Missegregation rates as determined in 1A and 4A, classified in euploid clones, clones harbouring whole chromosome aneuploidies, and clones harbouring segmental aneuploidies. Lines show the mean; error bars indicate standard deviation. C. Spearman correlation between the number of gained coding genes and the level of CIN as determined in A.

One of the segmental clones (SA6) with high CIN rates displayed aberrant sequence reads on the q-arm of chromosome 3 as observed by CNV sequencing (Fig. S2). We selected this clone for further characterization. We performed single-cell sequencing to determine the copy number variations per cell to get more insight into the variation between cells (Fig. 5A, B). Besides the expected gain of 10q, only few instabilities could be observed in the parental cells, in line with the live cell imaging data (Fig. 5A, Fig. 1A). Clone SA6 displayed the imbalances observed by CNV, namely a partial loss of chromosome 3 and 20, a loss of chromosome 13 and a gain of chromosome 14 and 19 (Fig. 5B). Strikingly, the extent of the loss of the terminal part of chromosome 3 was different in every single cell analyzed, a pattern also observed for the partial loss of chromosome 20 in a subset of the cells. All these observations were confirmed and further detailed by performing COBRA-FISH analysis that allows for the visualization of all different chromosomes (Fig. 5C-D). 95% of the cells displayed a derivative chromosome resulting from a translocation between 10 and 14. Also, we found in 15/40 (37,5%) analyzed metaphase cells that chromosome 3 was affected by translocations, dicentric chromosome formation or telomeric associations. The most frequent abnormality observed related to chromosome 3, also involved chromosome 20, which often formed a derivative dicentric chromosome at various breakpoints (Fig. 5A, D). Probing for specific locations of chromosome 3 showed that one copy of chromosome 3 had lost both signals representing the q-arm in clone SA6 (Fig 5E). The high frequency of dicentric chromosome formation along with ongoing loss of parts of chromosome 3 and 20 is consistent with an ongoing breakage-fusion-bridge (BFB)-cycle (Umbreit et al., 2020). We hypothesize that an initial missegregation led to the breakage of chromosome 3 and 20, and consequent fusion of the broken ends resulted in the formation of a dicentric chromosome, thereby triggering ongoing BFB-cycles. These data show that besides the general effects of genomic imbalances, segmental imbalances can instigate the formation of dicentric chromosomes as a result of fusion of the broken chromatin fragment with another chromosome and the consequent breakage-fusion-break cycles, which lead to CIN.

**Figure 5.**
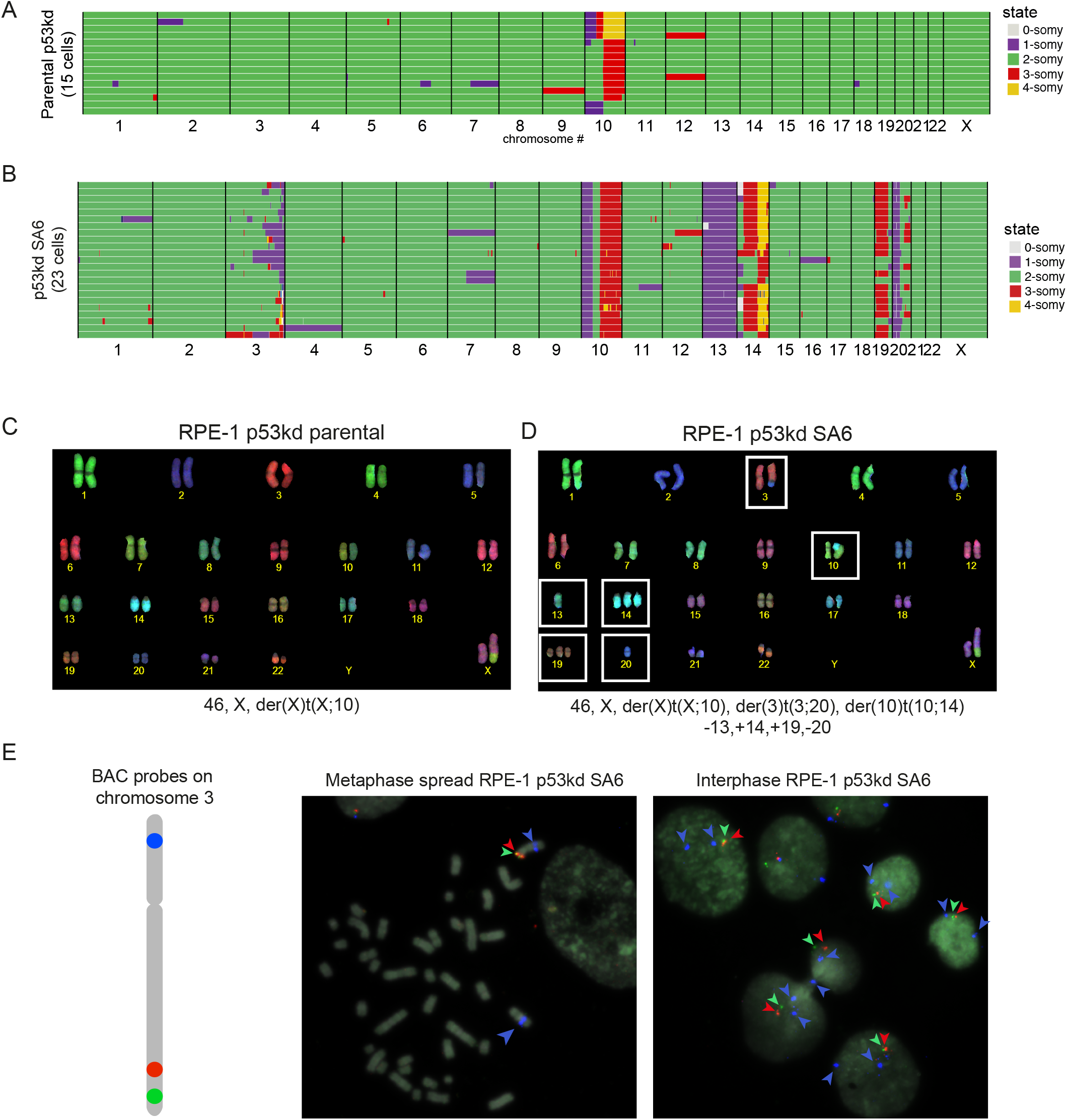
Segmental aneuploidies can lead to the onset of BFB-cycles via dicentric chromosome formation. A. Genome-wide chromosome copy number profiles of parental RPE-1 p53kd as determined by single-cell sequencing. Green indicates disomic chromosome regions, purple monosomic, red trisomic and yellow tetrasomic. Gain of the q-arm of chromosome 10 and occasional gain of chromosome 12 is expected. B. Genome-wide chromosome copy number profiles of clone SA6 as determined by single-cell sequencing as in A. Different patterns of aneuploidy are seen in chromosome 3. C-D. Representative images of chromosome spreads labelled with combined binary ratio labelling– fluorescence in situ hybridization of parental RPE-1 cells and clone SA6. The corresponding karyotypes are indicated below the images. White boxes indicate both numerical and segmental abnormalities specific to SA6. E) Metaphase (left) and interphase (right) representative images of cells from clone SA6, using BAC probes specific for different locations throughout the long arm (in red and green) and the short arm (in blue).

## Discussion

Aneuploidy and CIN are both prevalent features of tumors that highly correlate with each other (Ben-David and Amon, 2020). It has been proposed that aneuploidy can be a driver of CIN. Indeed, it has been shown that extra chromosomes can instigate a CIN phenotype (Sheltzer et al., 2011; Zhu et al., 2012; Duesberg et al., 1998; Nicholson et al., 2015; Passerini et al., 2016), however there are also examples where aneuploidy does not induce CIN (Lengauer et al., 1997; Sheltzer and Amon, 2011; Zhu et al., 2012). Moreover, there are cases where aneuploidy and CIN do not co-occur in tumors (Ben-David and Amon, 2020; Van Jaarsveld and Kops, 2016). This indicates that there must be specific aspects of aneuploidy that can contribute to CIN. To understand this, we set out to systematically investigate the impact of many different aneuploid karyotypes on CIN. While studies focusing on the consequences of chromosomal gains have been done extensively, research investigating the cellular consequences of monosomies or segmental aneuploidies have been limited to date. By creating aneuploid clones harboring many unique karyotypes, we were able to extensively evaluate the link between aneuploidy and CIN.

### Gene imbalances lead to impaired growth

In line with previous studies, we found that the cellular fitness of the vast majority of our aneuploid clones was decreased (Sheltzer et al., 2011, 2012; Stingele et al., 2012; Torres et al., 2007; Williams et al., 2008). We found a significant correlation between the number of imbalanced genes and reduced proliferation rates, and both the gains and losses of genes contributed to this correlation. In line with this, we found that both monosomic and trisomic clones were growth perturbed. We conclude that CIN is not a direct cause for slow growth as some fast-growing clones are highly unstable and some of the monosomic clones displayed a growth impairment although they are not CIN. This is in line with previous studies in yeast that found no correlation between CIN levels and growth impairment (Sheltzer et al., 2011; Zhu et al., 2012). It needs to be noted that the induction of acute CIN did cause a downregulation of cell cycle associated Hallmarks (Fig. 3D), which might indicate that CIN has a minor contribution to the slow growth phenotype. However, our findings indicate that the reduced growth of aneuploid cells is likely driven by the dosage changes of specific genes caused by both gains and losses and is not directly caused by the CIN phenotype.

### Monosomies are chromosomally stable

When evaluating clones harboring whole chromosome aneuploidies, we showed that aneuploidy *per se* is not always sufficient to induce CIN, as a subset of our clones were chromosomally stable. Remarkably, we found that clones that only harbor monosomies do not instigate a CIN phenotype while most clones harboring trisomies do. Importantly, the different CIN levels cannot simply be explained by a difference in karyotype complexity as trisomic clones with karyotype complexities similar to the monosomic clones do show increased CIN (for example compare WA3 to WA8). The different effects of monosomies and trisomies on CIN have not been studied extensively as most model systems studying the effects of aneuploidy on CIN only focused on chromosome gains (Nicholson et al., 2015; Passerini et al., 2016; Sheltzer et al., 2011; Zhu et al., 2012). However, recently it was documented that certain monosomies can trigger a CIN phenotype (Chunduri et al., 2021). There might be several explanations for this apparent discrepancy. First, it was reported that there are additional aneuploidies present in the background of a subset of the monosomy clones (for example a segmental gain of chromosome 22 in one clone). Such aneuploidies could contribute to the observed instability as segmental aneuploidies can instigate a CIN phenotype. Moreover, in the study by Chunduri et al, they observed chromatin bridges only in p53KO cells and not in monosomies generated in a p53kd background, that we used to generate our clones. Therefore, we cannot rule out that there is a role for minimal levels of p53 that are potentially left in our monosomic clones that would protect against CIN. However, this is unlikely as we do observe high CIN levels in the trisomic clones that were generated in the same background. Finally, since we only investigated a limited number of monosomic clones, it is possible that losing certain chromosomes can indeed lead to instability for instance due to certain haplo-insufficient genes involved in chromosome segregation that are present on these lost chromosomes.

### Trisomies cause CIN levels that correlate with the number of gained genes

Interestingly, we found a strong correlation between the levels of CIN and the amount of gained coding genes. It has been shown before in aneuploid yeast strains that the negative consequences of aneuploidy, under which genomic instability, are due to the presence of extra genes and not the presence of extra DNA (Sheltzer et al., 2011; Torres et al., 2007), suggesting that CIN is indeed induced by altered expression of genes and not simply a consequence of having extra chromosomes that need to participate in mitosis. However, these studies did not find a correlation between the amount of gained material and instability, suggesting that the underlying mechanisms driving CIN might differ between yeast and mammals.

So why do chromosome gains result in CIN and monosomies do not? Both gaining extra genetic material as well as losing genetic material will result in imbalances of proteins. Such expression changes will affect complex stoichiometry in specific cases. Cells deal with the excess of unincorporated proteins by performing dosage compensation (Brennan et al., 2019; Dephoure et al., 2014; Torres et al., 2007, 2010; Kojima and Cimini, 2019). This has been shown to occur both in trisomies (Stingele et al., 2012) as well as in monosomies (Chunduri et al., 2021). Although both monosomies and trisomies perform dosage compensation, the critical difference between monosomies and trisomies might be their distinct mechanism for dosage compensation and the associated stress pathways that are induced. Imbalances caused by monosomies could be resolved by upregulating the respective haploid gene products by decreasing protein turnover and/or increasing protein production while trisomies rather increase protein turnover by enhancing their degradation, often coinciding with enhanced levels of unfolded protein, ER stress and aggregate formation (Brennan et al., 2019; Dephoure et al., 2014; Santaguida and Amon, 2015; Torres et al., 2010; Zhang and Kaufman, 2008). In fact, studies in yeast indicate that the toxicity of individually overexpressed dosage-sensitive genes can be attributed by the enhanced burden on the protein turnover machinery (Makanae et al., 2013; Morrill and Amon, 2019). Indeed, interfering with protein folding and turnover in parental cells using low doses of inhibitors resulted in CIN, suggesting that interfering with the proteostasis machinery might be sufficient to drive chromosomal instability. Moreover, transcriptome analysis showed an activation of the PERK-branch of the UPR specifically in trisomies, suggesting ER stress to be selective for trisomic clones. However, how ER stress or saturation of the protein turnover machinery could drive CIN remains enigmatic. Future studies should shed light on the mechanisms of dosage compensation in trisomies and monosomies, if this indeed has a different impact on the stress pathways that are activated and how this can translate into CIN.

### Trisomies induce a stronger inflammatory response as compared to monosomies

The most prominent difference on transcriptome level between trisomies and monosomies, was the more significant upregulation of an inflammatory response in trisomies. As a similar response was observed in parental cells with induced acute aneuploidy, this suggests that this response is probably a consequence of the elevated levels of CIN in the trisomic clones. It is known that CIN can induce such a response via the cGAS-STING pathway, as a result of micronuclei rupture or chromatin bridges (Dou et al., 2017; MacKenzie et al., 2017) or via cGAS-independent activation of NF-kB (Wang et al., 2021). Possibly proteotoxic stress itself can contribute to an elevated inflammatory response as it has been shown that an overload of the ER with proteins accumulating inside the organelle can trigger an NF-kB response (Pahl and Baeuerle, 1995). We can also not exclude that the inflammatory response that is triggered specifically in trisomic cells, is contributing to the CIN phenotype, thereby generating a positive feedback that results in the maintenance of high CIN levels. More research is needed to reveal which exact aspects of CIN or which other factors are responsible for the elevated inflammatory response that we observe in trisomies.

### Segmental aneuploidies can result in BFB-cycles

Clones harboring segmental aneuploidies also display a link between the number of gained genes and instability. However, we found that segmental aneuploidies can have additional defects leading to chromosomal instability. By combining single cell sequencing and COBRA-FISH, we found strong evidence of an ongoing BFB-cycle, in at least one clone harboring segmental aneuploidies. This ongoing BFB-cycle was driven by a fusion between chromosome 3 and chromosome 20, resulting in a dicentric chromosome. Although BFB-cycles have been associated with telomere damage (Kinsella and Bafna, 2012), we show here that such segmental rearrangements can also arise after chromosome missegregation events resulting in broken chromosomes. Thus, faulty repair of segmental aneuploidies can result in the formation of abnormal derivative chromosomes, thereby leading to ongoing BFB-cycles, which can be an important mechanism for genomic amplifications seen in cancer (Saunders et al., 2000). If dicentric chromosome formation also occurs in the trisomic aneuploid clones that display CIN needs further investigation.

### Concluding remarks

Together, our findings show that aneuploidy *per se* does not induce chromosomal instability. We observed that clones harboring trisomies show various levels of CIN while monosomies are chromosomally stable. These elevated CIN levels correlate with a stronger activation of the inflammatory response in trisomies as compared to monosomies. Interestingly, levels of CIN correlate significantly with the amount of gained coding genes. Moreover, inhibiting protein folding or protein turnover pathways in parental cells is sufficient to induce CIN. We hypothesize that excess protein production is putting a burden on the protein turnover machinery and this results in CIN by a yet to be defined mechanism. Finally, we found that segmental aneuploidies can cause ongoing segregation errors by inducing BFB-cycles. This knowledge contributes to our understanding of the relationship between aneuploidy and CIN and how different types of aneuploidy can have different impacts on cancer initiation and development.

## Materials and Methods

### Cell culture, cell lines, and reagents

hTert-immortalized retinal pigment epithelium (RPE-1) cells were obtained from ATCC and RPE-1 p53kd cells were kindly provided by R. Beijersbergen. RPE-1 p53kd cells were generated by transduction with pRetroSuper-p53 (with the shRNA sequence 5’-CTACATGTGTAACAGTTCC-3’) and selected with Nutlin-3 for functional loss of p53. H2B-Dendra2 cells were made as described in (Soto et al., 2017). Cells described above were cultured at 37C at 5% CO2 in Advanced Dulbecco’s Modified Eagle Medium: Nutrient mixture F-12 (DMEM-F12) with Glutamax (GIBCO), supplemented with 12% FCS (Clontech), 100 U/ml penicillin (Invitrogen), 100 μg/ml streptomycin (Invitrogen) and 2mM UltraGlutamin (Lonza). Inhibitors were all dissolved in DMSO and were used at the following concentrations: GSK923295 50nM, NMS-P715 480nM, Nutlin3a 10μM, MG132 75nM and 150nM, 17-AAG 15nM and 30nM.

### Generating aneuploid clones

Clones were generated by treating RPE-1 p53kd cells with and Mps1 inhibitor (NMS-P715) and a CENP-E inhibitor (GSK923295) to induce whole chromosome aneuploidies and segmental aneuploidies as shown in Soto et al., 2017. A total of 2688 wells were examined for the presence of individual cells to ensure a single cell was present. Of those, 481 wells contained a single cell. To be confident that a missegregation took place, only wells containing a single cell harboring a micronucleus were selected, which were 119 cells. Out of these, 17 established until a full clone, while others stopped proliferating. CNV sequencing showed an aneuploid karyotype for 13 clones. Thus, out of 481 single plated cells, we obtained 13 aneuploid clones, a success percentage of 3,5% Live cell imaging of these clones showed that aneuploid clones grew slower. To also generate clones that did not experience a MN during their generation, we selected established clones with a slow growth phenotype. With this approach we significantly improved the success rate to 17% (out of all single cell plated cells). These are the clones starting from WA8, and clone WA5.

The gain of 10q in parental RPE-1 cells, deriving from an imbalanced fusion of the q-arm of chromosomes 10 to the X chromosome ((Janssen et al., 2011), ATCC) and chromosome 12, present in a fraction of RPE-1 cells (Soto et al., 2017; Zhang et al., 2015) were not considered *de novo* aneuploidies (Fig. S1).

### Time-lapse live-cell imaging

For live-cell imaging, cells were grown in Lab-Tek II chambered coverglass (Thermo Science). Images were acquired every 5 minutes using a DeltaVision Elite (Applied Precision) microscope maintained at 37C, 40% humidity and 5% CO2, using a 20x 0.75 NA lens (Olympus) and a Coolsnap HQ2 camera (Photometrics) with 2 times binning. Image analysis was done using ImageJ software. DNA was visualized using 0,25uM siR-DNA (Spirochrome).

### Cell growth analysis

Proliferation was measured by using the Incucyte FRL (Essen BioScience) or the Lionheart FX automated microscope (Biotek). For the Incucyte, 250 cells were plated in 96-well plates. Three or four replicate wells were imaged per cell line (phase-contrast) with a 4 h interval for 6 d. Confluency was determined by IncuCyte FLR software 2011A Rev2 and IncuCyte Zoom software 2013B Rev1 using phase-contrast images, and doubling times were calculated using GraphPad Prism 6 software.

For the Lionheart, 500 cells were plated in 96-well plates. Two or three replicate wells were imaged per cell line with a 4 h interval for 5 days, stained with DNA dye siR-DNA (Spirochrome). Proliferation rates were measured by performing cell count analysis using Gen5 software (BioTek) and doubling times were calculated using GraphPad Prism 8 software.

### Copy number analysis

DNA was isolated using the DNeasy Blood and Tissue kit (Qiagen) according to the manufacturer’s protocol. The amount of double-stranded DNA in the genomic DNA samples was quantified by using the Qubit. dsDNA HS Assay Kit (Invitrogen, cat no Q32851). Up to 2000 ng of double-stranded genomic DNA was fragmented by Covaris shearing to obtain fragment sizes of 160-180 bp. Samples were purified using 1.8X Agencourt AMPure XP PCR Purification beads according to the manufacturer’s instructions (Beckman Coulter, cat no A63881). The sheared DNA samples were quantified and qualified on a BioAnalyzer system using the DNA7500 assay kit (Agilent Technologies cat no. 5067-1506). With an input of maximum, 1 μg sheared DNA, library preparation for Illumina sequencing was performed using the KAPA HTP Library Preparation Kit (KAPA Biosystems, KK8234). During library enrichment, 4-6 PCR cycles were used to obtain enough yield for sequencing. After library preparation, the libraries were cleaned up using 1X AMPure XP beads. All DNA libraries were analyzed on a BioAnalyzer system using the DNA7500 chips for determining the molarity. Up to eleven uniquely indexed samples were mixed together by equimolar pooling, in a final concentration of 10nM, and subjected to sequencing on an Illumina HiSeq2500 machine in one lane of a single read 65 bp run, according to manufacturer’s instructions.

### Calculation of number of imbalanced/gained/lost genes

For clones harboring whole chromosome aneuploidies, the number of imbalanced/ gained / lost coding genes was calculated by determining the aneuploid chromosomes per clone using CNVseq data. Then per clone the amount of coding genes located on the monosomic chromosomes were summed up to obtain amount of lost coding genes, the amount of coding genes located on the trisomic chromosomes were summed up to obtain amount of gained coding genes and for the number of imbalanced genes, the total of lost and gained coding genes was determined. To determine the amount of coding genes per chromosome we made use of Ensembl release 79 using the genome assembly GRCh38, only considering protein-coding genes. For segmental aneuploidies we determined the breakpoints using CNVseq data and calculated the amount of coding genes on the gained, lost or imbalanced segments with the same method as described above.

### Single-cell whole genome sequencing (scWGS), data processing and analysis

Single cells were lysed and stained in a nuclei isolation buffer (100 mM Tris-HCl [pH 7.4], 150 mM NaCl, 1 mM CaCl2, 0.5 mM MgCl2, 0.1% NP-40, and 2% BSA). Nuclei were stained with propidium iodide and Hoechst 33258 at concentrations of 10 μg/mL. Individual nuclei of G1 cells were sorted directly into 5 μL freezing buffer (50% PBS, 7.5% DMSO, and 42.5% 2X ProFreeze-CDM [Lonza]) in 96-well plates using a FACSJazz cell sorter (BD Biosciences). Plates were spun down at 500g for 5 min at 4C prior to storage at -80C until library preparation. Pre-amplification free scWGS library preparation was performed as described previously (van den Bos et al., 2016). Libraries were sequenced on an Illumina HiSeq2500 sequencing platform, with clusters generated using the cBot. Raw sequencing data were demultiplexed and converted into fastq format using standard Illumina software (bcl2fastq version 1.8.4). Indexed bam-files were generated by mapping to GRCh37 using bowtie2 (version 2.2.4). Duplicate reads were marked using BamUtil (version 1.0.3). Copy number variations were called using the R-package AneuFinder, and quality control were performed as described before (Bakker et al., 2016). The analysis was done using 1Mb bins.

### COBRA-FISH analysis

From RPE-1 p53kd parental and clone SA6, cells were harvested using a metaphase harvesting protocol described earlier (Janssen et al., 2011). Metaphase cells were further analyzed using a multicolor-FISH karyotyping technique called COBRA-FISH, allowing identification of chromosomes based on spectrally distinct colors, according to protocols described earlier (Szuhai and Tanke, 2006).

### Immunoblotting

RPE-1 cells were harvested and lysed using Laemmli buffer (120 mM Tris, pH 6.8, 4% SDS, and 20% glycerol). Equal amounts of protein were separated on a polyacrylamide gel and subsequently transferred to nitrocellulose membranes. Membranes were probed with the following primary antibodies: LC3B (Rabbit, Sigma-Aldrich, L7543), alfa-Tubulin (mouse, Sigma, t5168). HRP-coupled secondary antibodies (Dako) were used in a 1:1000 dilution. The immunopositive bands were visualized using ECL Western blotting reagent (GE Healthcare) and a ChemiDoc MP System (Biorad).

### RNA sequencing and data analysis

RPE-1 p53KD cells (parental and clones) were harvested in buffer RLT (Qiagen). Strand-specific libraries were generated using the TruSeq PolyA Stranded mRNA sample preparation kit (Illumina). In brief, polyadenylated RNA was purified using oligo-dT beads. Following purification, the RNA was fragmented, random-primed and reserve transcribed using SuperScriptII Reverse Transcriptase (Invitrogen). The generated cDNA was 3′ end-adenylated and ligated to Illumina Paired-end sequencing adapters and amplified by PCR using HiSeq SR Cluster Kit v4 cBot (Illumina). Libraries were analyzed on a 2100 Bioanalyzer (Agilent) and subsequently sequenced on a HiSeq2000 (Illumina). We performed RNAseq alignment using TopHat 2.1.1. on GRCh38 and counted reads using Rsubread 2.4.3 (Ensembl 102). We calculated differential expression between two biological replicates of the parental and each clone, between the monosomic (WA1, WA2, WA5) and trisomic clones (WA8, WA11, WA12), as well as the Mps1 and CENP-E inhibitor treated vs control, using DESeq2 1.31.3. We tested for gene set differences by using a linear regression model of the Wald statistic (as reported by DESeq2) between genes belonging to a set vs. genes not belonging to a set. Gene set collections included MSigDB Hallmarks (2020) and Gene Ontology (2021). For the UPR focus, we chose genes annotated with GO categories GO:0036500 (ATF6-mediated unfolded protein response), GO:0036498 (IRE1-mediated unfolded protein response), and GO:0036499 (PERK-mediated unfolded protein response) or their child terms.

### Growth assay to determine IC50

250 cells were plated in 96-well plates (BD Biosciences) and drugs were added on day 1 using a Digital Dispenser (Tecan Männedorf). On day 8, cells were fixed for 10 minutes in 99% methanol and stained with 0,1% crystal violet. After 4 hours, staining solution was removed and plates were washed 4 times with water after which plates were air dried. Plates were scanned and analysed with ImageJ software (NIH) and relative cell survival plots were generated and IC50 was calculated with Prism 7 (GraphPad).

## Acknowledgements

We would like to thank the Medema, Rowland and Jacobs lab for helpful discussions. This study was supported by funds from the Dutch Cancer Society (KWF-Young Investigator Grant-12233) granted to J.A.R.. We thank the Genomics Core Facility of the Netherlands Cancer Institute for sample preparation, data acquisition, and analysis of CNV and RNA sequencing experiments.

## Author contributions

R.H.M., M.S and J.A.R. conceived the project. D.C.H, M.S., K.S. and J.A.R., B.B., and D.C.J.S. performed and analyzed the experiments. M.Sch performed data processing and data analysis. B.B. analyzed the single-cell sequencing data. F.F. and P.M.L. provided resources and input on the project. D. C. H, M.S., J.A.R., and R.H.M. wrote the manuscript.

## Supplementary Figure Legends

**Supplementary Figure 1.**
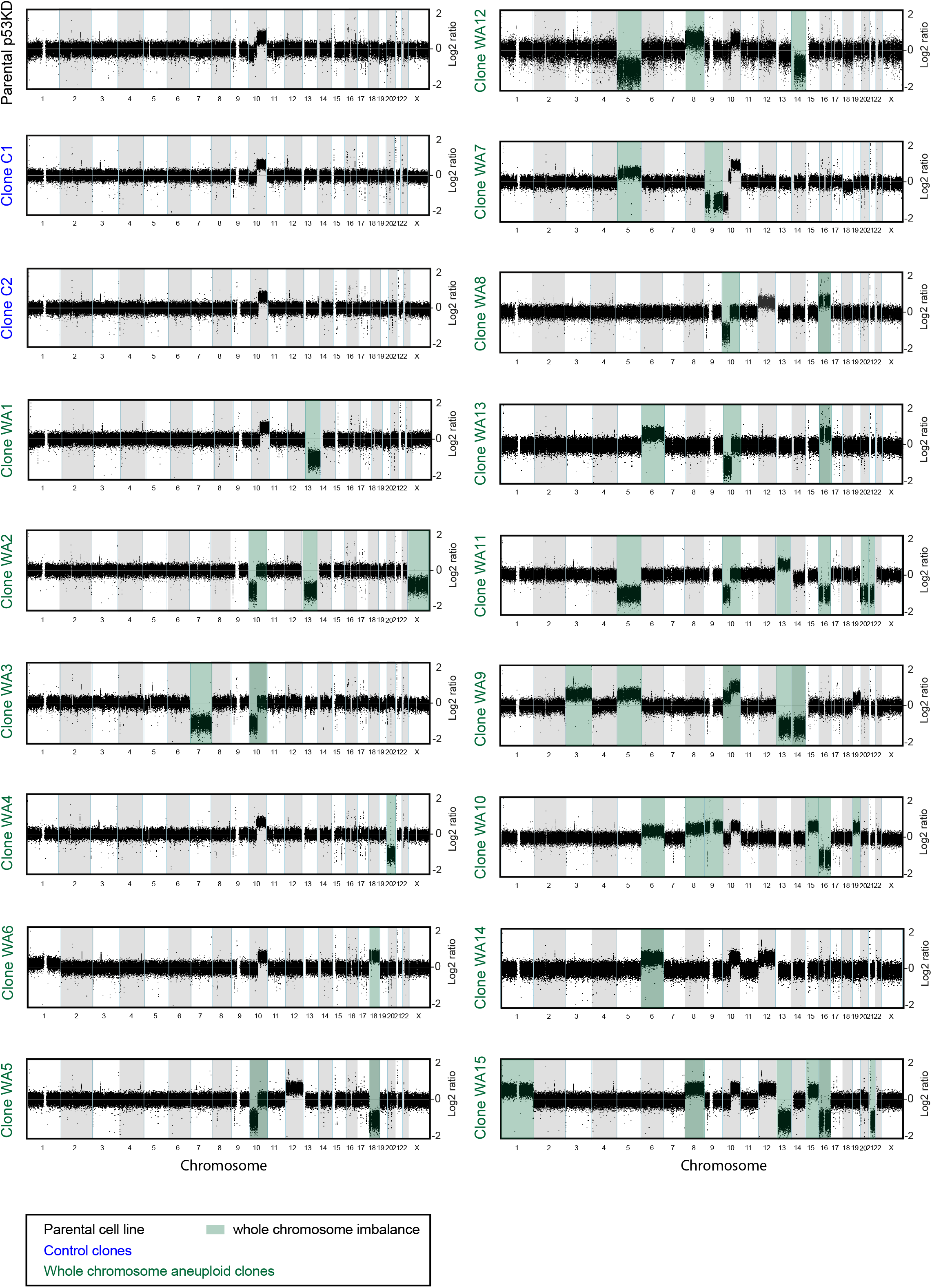
Characterization of euploid clones and whole chromosome aneuploid clones. Genome-wide chromosome copy number profile as determined by CNV-seq of the RPE-1 p53kd parental clone (labelled in black), two euploid clones (labelled in blue) and 10 clones harboring solely whole chromosome imbalances (labelled in green). Chromosome gains and losses were depicted in green boxes. Alterations of chromosome 10 and 12, already present in the parental cells, were not highlighted.

**Supplementary Figure 2.**
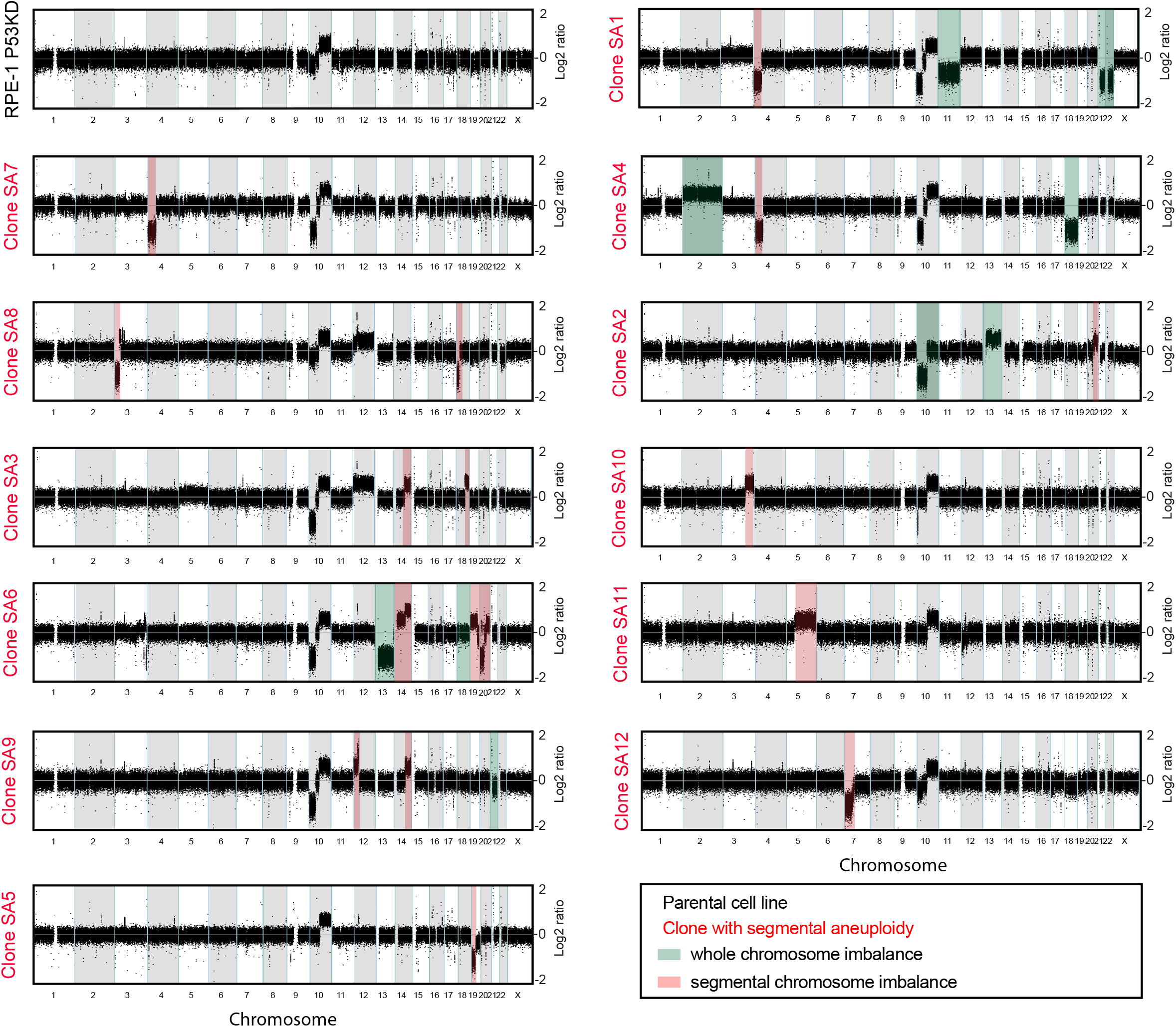
Characterization of segmental chromosome aneuploid clones. Genome-wide chromosome copy number profile as determined by CNV-seq of the RPE-1 p53kd parental clone (labelled in black), two euploid clones (labelled in blue) and 10 clones harboring segmental chromosome imbalances (labelled in red). Chromosome gains and losses were depicted in green boxes. Alterations of chromosome 10 and 12, already present in the parental cells, were not highlighted.

**Supplementary Figure 3.**
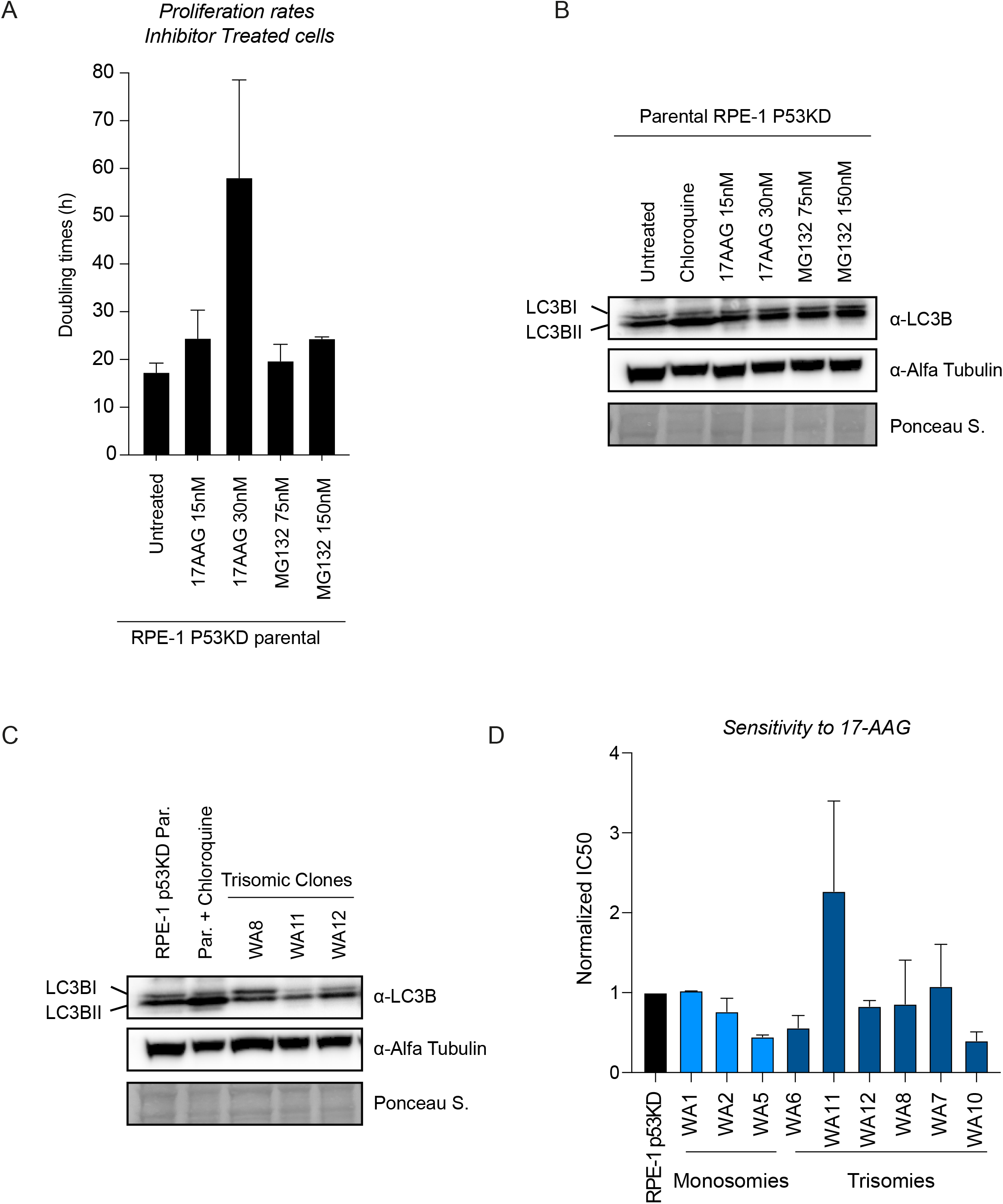
No detectable proteotoxic stress in trisomies or in parental cells treated with proteostasis interfering drugs. A. Doubling time of parental p53KD cells untreated and treated with different concentrations of proteostasis interfering drugs, as measured in Fig. 2. B. Immunoblot showing conversion from LC3B-I to LC3B-II in parental p53KD cells untreated, treated with 50uM chloroquine and treated with low doses of proteostasis interfering drugs for 24 hours. Loading control is Alpha Tubulin. C. Immunoblot showing conversion from LC3B-I to LC3B-II in parental p53KD cells untreated, treated with 50uM chloroquine to block autophagy as a positive control and 3 different trisomic clones. Loading control is Alpha Tubulin. D. IC50 values of Hsp90 inhibitor 17-AAG as determined by growth assays of parental p53KD cells and various trisomic clones, ordered per amount of gained coding genes.

**Supplementary Figure 4.**
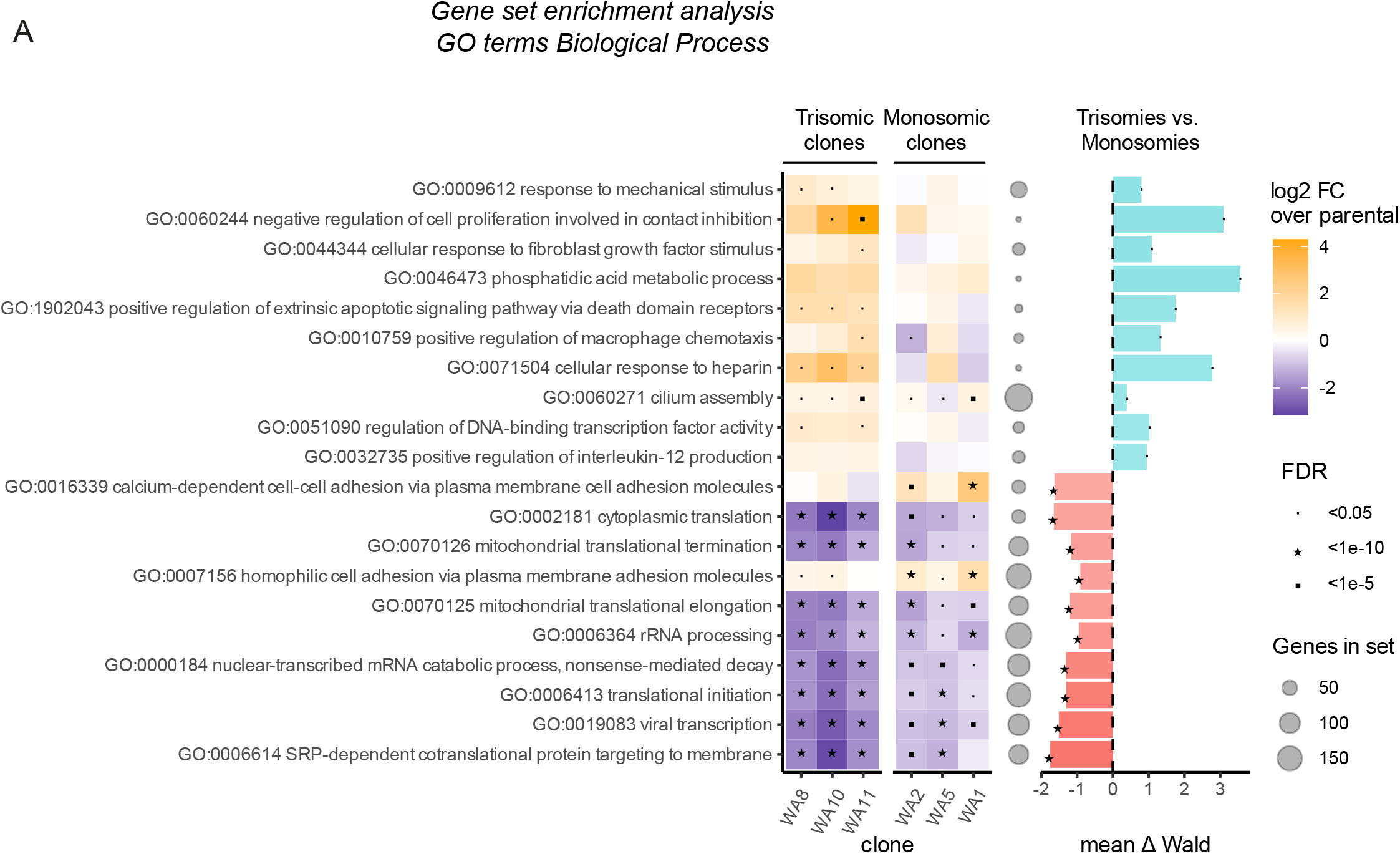
Differentially expressed GO Biological processes between Trisomic and Monosomic clones. A. Gene set enrichment analysis (GSEA) of RNA sequencing data, evaluating up and downregulated GO Biological Processes in trisomy clones and monosomy clones compared to parental (left graph). Two replicates of every clone were sequenced and the Log2 fold change (FC) was determined compared to parental cells. The false discovery rates (FDR) are indicate with symbols. Largest differences between trisomies and monosomies are shown on the right. Differences in Hallmarks between trisomies and monosomies were determined by Wald statistical testing.

